# Genetic architecture of inbreeding depression may explain its persistence in a population of wild red deer

**DOI:** 10.1101/2023.11.23.568439

**Authors:** Anna M. Hewett, Susan E. Johnston, Josephine M. Pemberton

## Abstract

Inbreeding depression is of major concern in declining populations, but relatively little is known about its genetic architecture in wild populations, such as the degree to which it is composed of large or small effect loci and their distribution throughout the genome. We combine fitness and genomic data from a wild population of red deer to investigate the genomic distribution of inbreeding effects. Inspired by the runs of homozygosity (ROH)-based inbreeding coefficient, F_ROH_, we use chromosome-specific inbreeding coefficients (F_ROHChr_) to explore whether the effect of inbreeding varies between chromosomes. Under the assumption that within an individual the probability of being identical-by-descent is equal across all chromosomes, we used a multi-membership model to estimate the deviation of F_ROHChr_ from the average inbreeding effect. This novel approach ensures effect sizes are not overestimated whilst maximising the power of our available dataset containing >35,000 autosomal SNPs. We find that most chromosomes confer a minor reduction in fitness-related traits, which when these effects are summed, results in the observed inbreeding depression in birth weight, survival and lifetime breeding success. However, no chromosomes had a significantly detrimental effect compared to the overall effect of inbreeding. We conclude that in this population, inbreeding depression is the result of multiple mild or moderately deleterious mutations spread across all chromosomes. As predicted by genetic theory these mutations will be inefficiently purged, explaining the persistence of inbreeding depression in this population.

## Introduction

The reduced fitness of inbred individuals, known as inbreeding depression, is of particular concern for the management and conservation of declining species. Accumulation of inbreeding depression effects can lead to reduced population growth and in the worst cases extinction (Hedrick and Kalinowski, 2000; Keller and Waller, 2002). A number of wild populations across a range of taxa have been reported to experience inbreeding depression in various fitness-related traits including fecundity, survival, body weight, germination success and susceptibility to disease. (e.g. Crnokrak and Roff, 1999; Hajduk *et al*, 2018; Keller and Waller, 2002; Liberg *et al*, 2005). Moreover, in the wild the overall cost of inbreeding depression is likely higher in natural populations than for those in captivity (Ralls *et al*, 1988). Along with the increased risk of extinction, natural populations are more affected by environmental factors such as extreme weather, food scarcity or disease all of which will influence an individual’s probability of survival (Crnokrak and Roff, 1999).

The two main hurdles to overcome in order to estimate inbreeding depression in natural populations are obtaining measures of fitness and obtaining a detailed pedigree from which to estimate meaningful individual inbreeding coefficients (Pemberton, 2008). Advances in genomic technologies, such as SNP-arrays and whole genome sequencing, have helped overcome the latter issue, instead enabling more precise marker-based estimations of inbreeding (Hedrick and Garcia-Dorado, 2016; Kardos *et al*, 2015). A handful of studies of non-model organisms living in the wild have since shown evidence for inbreeding depression using genomic data (e.g. Bérénos *et al*, 2016; Hoffman *et al*, 2014; Huisman *et al*, 2016; Kardos *et al*, 2023; Khan *et al*, 2021; Stoffel *et al*, 2021). Suitable data from wild populations can still be hard to obtain, particularly for large samples of individuals, but rising affordability of genomic data should increase the prevalence of these types of studies (Kardos *et al*, 2016). The unique value of genomic studies is the ability uncover information on the genetic architecture of inbreeding depression, such as identifying loci with large effects and determining the genomic distribution of effects (Hedrick and Garcia-Dorado, 2016).

Genomic data may further uncover the extent to which recessive deleterious alleles or overdominant loci contribute to inbreeding depression in wild populations, which is still largely unknown (Kardos *et al*, 2016). Inbreeding increases homozygosity in the population, which, in both cases, leads to an increased frequency of the less fit genotype, either due to expressing the deleterious effect or due to not expressing the heterozygote phenotype, respectively. Experimental evidence supports the existence of both these mechanisms but implies that overdominance is rare and of lesser importance to inbreeding depression than recessive deleterious mutations (Charlesworth and Willis, 2009; Falconer and Mackay, 1996). Slightly counterintuitively, severe inbreeding can reduce inbreeding depression by facilitating purifying selection, purging deleterious recessive mutations from the population. Theory shows that purging is faster and more effective for large-effect mutations as selection is stronger against these mutations, whereas the effects of inbreeding depression that are the result of multiple recessive small effect loci will be less easily purged (Charlesworth and Willis, 2009). Moreover, the rate of inbreeding also affects the efficiency of purifying selection. Under fast inbreeding (i.e. high ΔF), selection is less efficient than if inbreeding is slower (lower ΔF) (Wang *et al*, 1999).

In our study population of red deer inhabiting the Isle of Rum inbreeding depression has previously been demonstrated in several fitness-related traits (Coulson *et al*, 1999; Coulson *et al*, 1998; Slate *et al*, 2000; Walling *et al*, 2011), most recently by Huisman *et al* (2016) who used individual inbreeding coefficients estimated from a SNP-derived genomic relatedness matrix. Huisman *et al* (2016) showed inbreeding depression in birth weight, juvenile survival and lifetime breeding success (LBS) in both sexes. For an individual which is a product of a half sibling mating (F≈0.125) LBS was decreased by 95% and 72% in males and females respectively (Huisman *et al*, 2016). Since this study, we have obtained precise genomic locations of SNPs through an assembly of the red deer (*Cervus elaphus*) genome (Pemberton *et al*, 2021),allowing for the identification of runs of homozygosity (ROH). ROH are a direct consequence of identity-by-descent in the genome and, depending on the density of genomic markers available, can allow for fine-scale measures of inbreeding and localisation of causal loci. For example, for a simple monogenic disease, a causal recessive mutation can be identified based on presence or absence of disease in relation to segments of ROH (Kijas, 2013). More recently ROHs have been used to identify genomic regions containing large effect loci in a range of complex traits (Hill *et al*, 2022; Pryce *et al*, 2014; Stoffel *et al*, 2021), highlighting the value of using ROHs in this type of analysis. Here, given the density of SNP genotypes available, our aim was to use ROH to identify whether specific chromosomes underpin inbreeding depression in this wild population of red deer. Using a modified version of the ROH-based inbreeding coefficient (F_ROH_) we quantified chromosome-specific inbreeding coefficients (F_ROHChr_), and then tested whether any chromosomes disproportionately contribute to inbreeding depression.

## Methods

### Genotyping and ROH calling

DNA was extracted from ear punches, *post mortem* tissues and cast antlers (see Huisman *et al* (2016) for details) for 3,198 individuals living within the red deer study population on the Isle of Rum between 1958 and 2022. Genotyping of DNA samples was performed on the Cervine Illumina 50K BeadChip (Brauning, 2015). SNP genotypes were clustered and scored using Illumina GenomeStudio v2.0 and the following quality control was imposed: SNP minor allele frequency (MAF) > 0.001, ID genotyping success >0.9, and SNP genotyping success >0.99; leading to 39,587 SNPs being retained at this stage. SNPs were mapped to the red deer (*Cervus elaphus*) genome assembly mCerEla1.1 (Pemberton *et al*, 2021).

We used the –homozyg function in *Plink* v2.0 to search for ROH in the population. The following parameters were used to identify ROH using a 35 SNP sliding window: 40 SNPs as the minimum number of SNPs in a ROH, minimum length of a ROH 2500 kb, minimum density of 1 SNP per 70 kb, 4 missing SNPs allowed in a window, 0 heterozygote SNPs allowed in a ROH, and a minor allele frequency threshold of 0.01. 36,997 mapped SNPs passed automatic filter and quality controls imposed by the *Plink* parameters used, all individuals passed quality controls.

### Chromosome-specific inbreeding coefficient

Chromosome-specific inbreeding coefficients (F_ROHchr_) for each individual were analogous to the genome-wide inbreeding coefficient, F_ROH_, where F_ROH_ is the sum of Mb in a ROH divided by the total autosome size. For every individual we summed the total Mb in a ROH on each autosome, then divided this by the corresponding chromosome length in Mb:

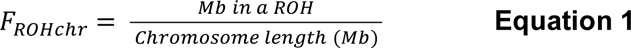

Chromosome lengths were obtained from the red deer genome assembly, mCerEla1.1 (Pemberton *et al*, 2021).

### Study traits

#### Estimated birth weight (kg)

We used neonatal capture weight in kg as a proxy for calf weight at birth. Calves are sampled and weighed soon after birth, their age in hours is also estimated based on field observations. In this model we fitted both the mother’s ID and an inverse pedigree relatedness matrix to account for known maternal effects and additive genetic variance (Gauzere *et al*, 2020), see Table 1 for additional covariates.

**Table 1:**
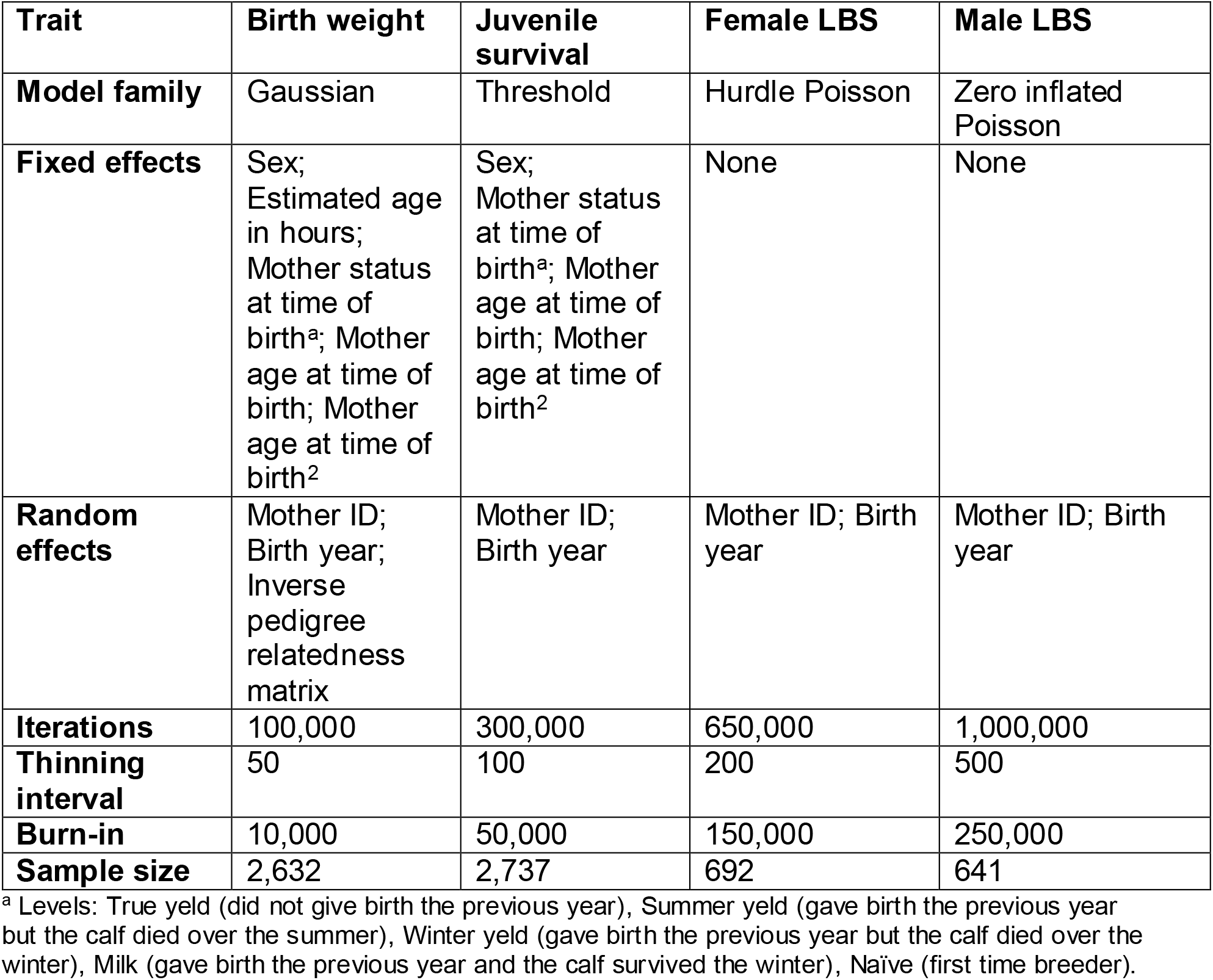
Model details for the investigated traits. All models were run in MCMCglmm. For each trait column, rows show the model family type; fixed effects; variables treated as random effects; number of iterations the model was run for; thinning interval (i.e. the intervals at which the Markov chain is stored); the burn-in and the number of individuals in each input data frame.

#### Juvenile survival (age 0-2 years)

Juvenile survival is considered as a binary response variable i.e. either 0 or 1. Any individual that died before the month of June two years after the year of birth was assigned a juvenile survival value of 0. Any individual that survived this time period was assigned a juvenile survival of 1. For most individuals an accurate death date is known from regular censuses throughout the year and mortality searching in winter, allowing for easy calculation of juvenile survival. For individuals with no known death information an estimated death year was assigned when the individual had not been sighted for three years. Individuals that died due accidents, desertion following tagging or being shot after ranging outside the study area before 2 years of age were removed from the dataset. Individuals that were shot as adults when ranging outside the study area were recorded as having a juvenile survival of 1.

For this and all subsequent traits no inverse relatedness matrix was fitted as previous studies show they have low heritability (Gauzere *et al*, 2020), Since there is inbreeding depression in birth weight, during modelling of juvenile survival we ran models with and without estimated birth weight (kg) fitted as a fixed effect to check the extent to which any inbreeding depression in juvenile survival was due to inbreeding depression in birth weight. This had no effect on the mean posterior estimates of F_ROHchr_ (though it did widen the confidence intervals, see Supplementary Figure 3). The final model presented does not include birth weight since it appears any inbreeding depression in birth weight is subsumed within inbreeding depression in juvenile survival.

#### Female lifetime breeding success (LBS)

Total female LBS was calculated for females that were born before 2006 that died of natural causes or were still alive in 2023 (n=10), so near the end of their reproductive span. Female breeding success is known from close field observation confirmed by pedigree reconstruction (Huisman 2016). We used a hurdle Poisson process to model female LBS as if they reach reproductive age, most females will reproduce at least once during their lifetime. In this case the ‘hurdle’ denotes survival to reproductive age and the truncated Poisson distribution denotes individuals with LBS>0.

#### Male lifetime breeding success (LBS)

Total LBS was calculated for all males that rutted in the study area and died of natural causes. We omitted records for all individuals born from 2008 onwards to avoid biasing results towards males that die young. Males can continue rutting until ∼15 years of age, hence, any male born after 2008 may not have reached their total LBS. Paternity is assigned through pedigree reconstruction using SNP genotypes and the pedigree inference software SEQUIOA (Huisman 2017). We used a zero-inflated Poisson process to model male LBS in the knowledge that a male can have zero LBS as a result of two causes: death before reproductive age (represented through the zero inflation) and failure to breed successfully (captured in the Poisson sampling distribution).

### Statistical analysis of chromosome-specific inbreeding effects

Our aim was to determine whether inbreeding depression is evenly distributed across chromosomes. We focussed on four traits outlined above with evidence for inbreeding depression identified from previous studies. We used a multi-membership approach to explore whether the effect of inbreeding varies between chromosomes for each trait, under the assumption that within individuals the F_ROHchr_ values are expected to be related as inbreeding is genome-wide. In this approach, the effect of each F_ROHchr_ is estimated as a deviation from the average effect of inbreeding. For each individual, inbreeding coefficients were summed across all chromosomes, which when fitted as a fixed effect describes the average effect of inbreeding among chromosomes. By treating all F_ROHchr_ as random effects and estimating their deviation this ensured we did not overestimate the effect of individual F_ROHchrs_.

For each trait the 𝑖^𝑡ℎ^individual was modelled as:

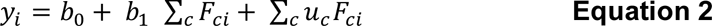

Where, 𝑏_0_ is the intercept and 𝐹_𝑐𝑖_ is the inbreeding coefficient for an individual 𝑖 on chromosome 𝑐. 𝑏_1_ is the estimated average effect of inbreeding and 𝑢_𝑐_ is the deviation from the average effect for chromosome 𝑐. Trace plots were inspected for validation of model convergence. Specific model parameters including additional fixed and random effects for each trait are shown in Table 1.

We used the 95% confidence intervals of the posterior estimates to assess the significance of each F_ROHchr._ Chromosomes where confidence intervals did not overlap the average effect of inbreeding were deemed significantly different. To calculate absolute F_ROHchr_ effects for each chromosome for trait predictions we combined each estimated deviation of F_ROHchr_ with the average effect of inbreeding. Trait predictions where inbreeding occurs equally on all chromosomes assumes that F_ROHchr_ is the same on all chromosomes, effects are summed to get the overall effect of inbreeding on each trait. Trait predictions where chromosomes are treated as independent assume that the inbreeding coefficient is present only on the focal chromosome and all other chromosomes are outbred (i.e. F_ROHchr_ = 0). All statistical models were run in the MCMCglmm R package (Hadfield, 2010).

## Results

The mean chromosome-specific inbreeding coefficient (F_ROHchr_) was 0.064 and ranged between 0 and 0.94, with the highest F_ROHchr_ found in two individuals on chromosome 3 (see outliers in Supplementary Figure 1). F_ROHchr_ within individuals were generally weakly positively correlated, except for chromosomes 2 and 21, 7 and 13, 12 and 18, and 31 and 14 which were slightly negatively correlated, see Figure 1.

**Figure 1.**
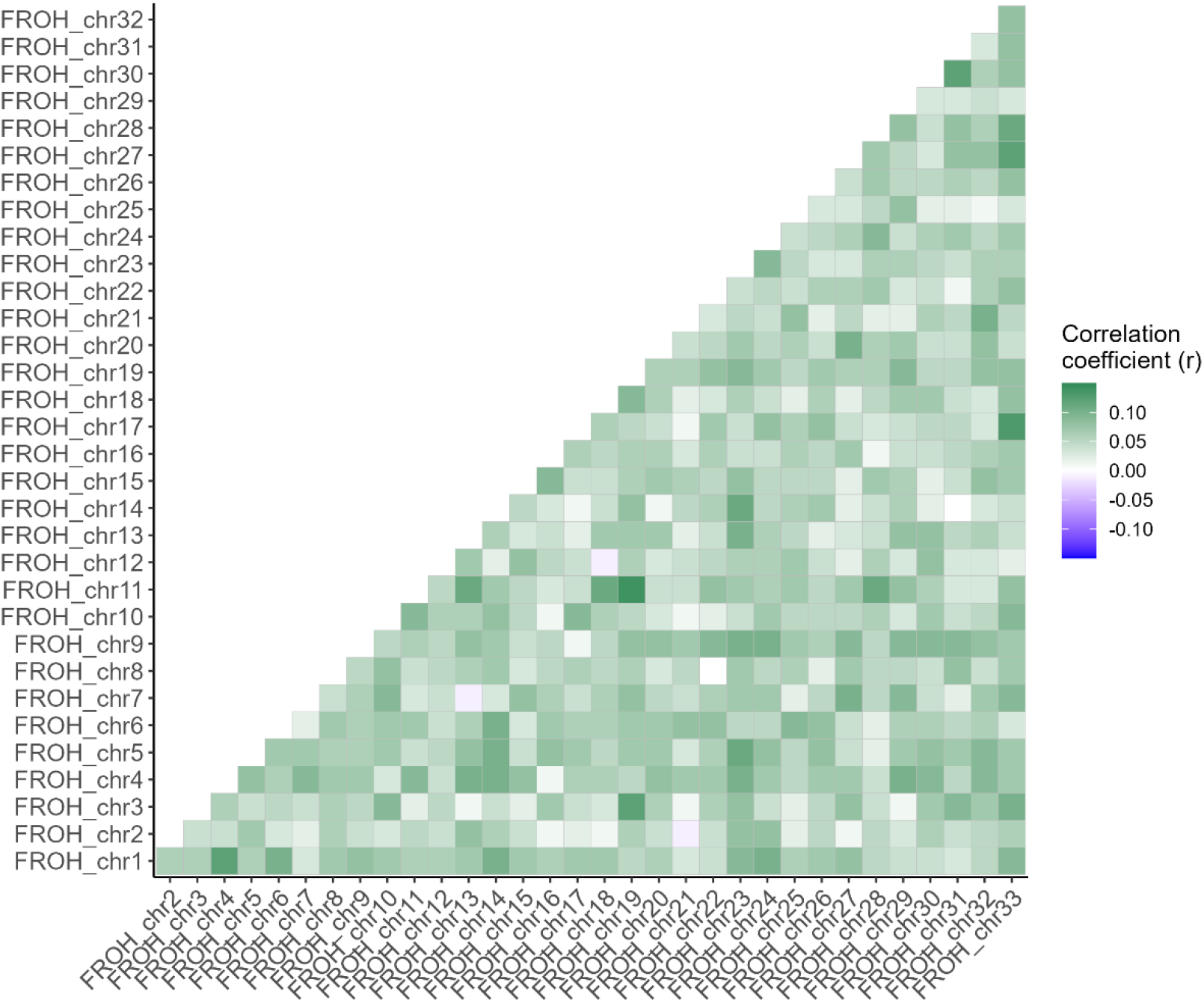
Correlation matrix of all chromosome-specific inbreeding coefficients (F_ROHchr_) within individuals. Green shows a positive correlation between F_ROHchr_, with stronger correlations shown in darker green, up to a maximum Pearson’s correlation coefficient of 0.15. Negative correlations are shown in purple, with darker purple showing stronger negative correlations up to a minimum of Pearson’s correlation coefficient of −0.15.

### Evidence for inbreeding depression

Overall, higher inbreeding coefficients were associated with significantly lower birth weights and probability of survival to age 2 (P-values <0.001). A sex-averaged calf with a F_ROHchr_ of 0.25 on all chromosomes was predicted to weigh 6.14 kg (CIs: 5.83 - 6.43kg) at birth, whereas, a completely outbred individual (F=0) was predicted to weigh 7.08kg (CIs: 6.91 - 7.25kg), see Fig 2A. For the probability of survival, the equivalent predictions were 36% (CIs: 21 – 54%) and 74% (CIs: 69 – 80%), Fig. 2B. We found a significant effect of inbreeding coefficient on whether a female ‘crossed the hurdle’ (i.e. had at least 1 offspring, p-value: 0.017), but no effect on the truncated Poisson process (number of offspring given the female had at least one offspring, p-value: 0.983). The combination of these processes predicts that a female with an inbreeding coefficient of 0.25 on all chromosomes has a LBS of 1.8 (CIs: 0.6 – 3.9) compared to a completely outbred female which has a predicted LBS of 4.6 (CIs: 3.6 – 5.6) (Fig 2C). In males we found a marginally significant association of overall inbreeding coefficient with the zero inflation element (p-value: 0.048) and a stronger association with the Poisson process (p-value<0.001). The combined effect of both processes shows a male with an inbreeding coefficient of just 0.1 is barely predicted to have one offspring, LBS prediction of 0.8 (CIs: 1.8 – 0.3) whereas a completely outbred male is predicted to have a total of 11.4 (CIs: 19.4 – 6.2) offspring (Fig 2D).

**Figure 2.**
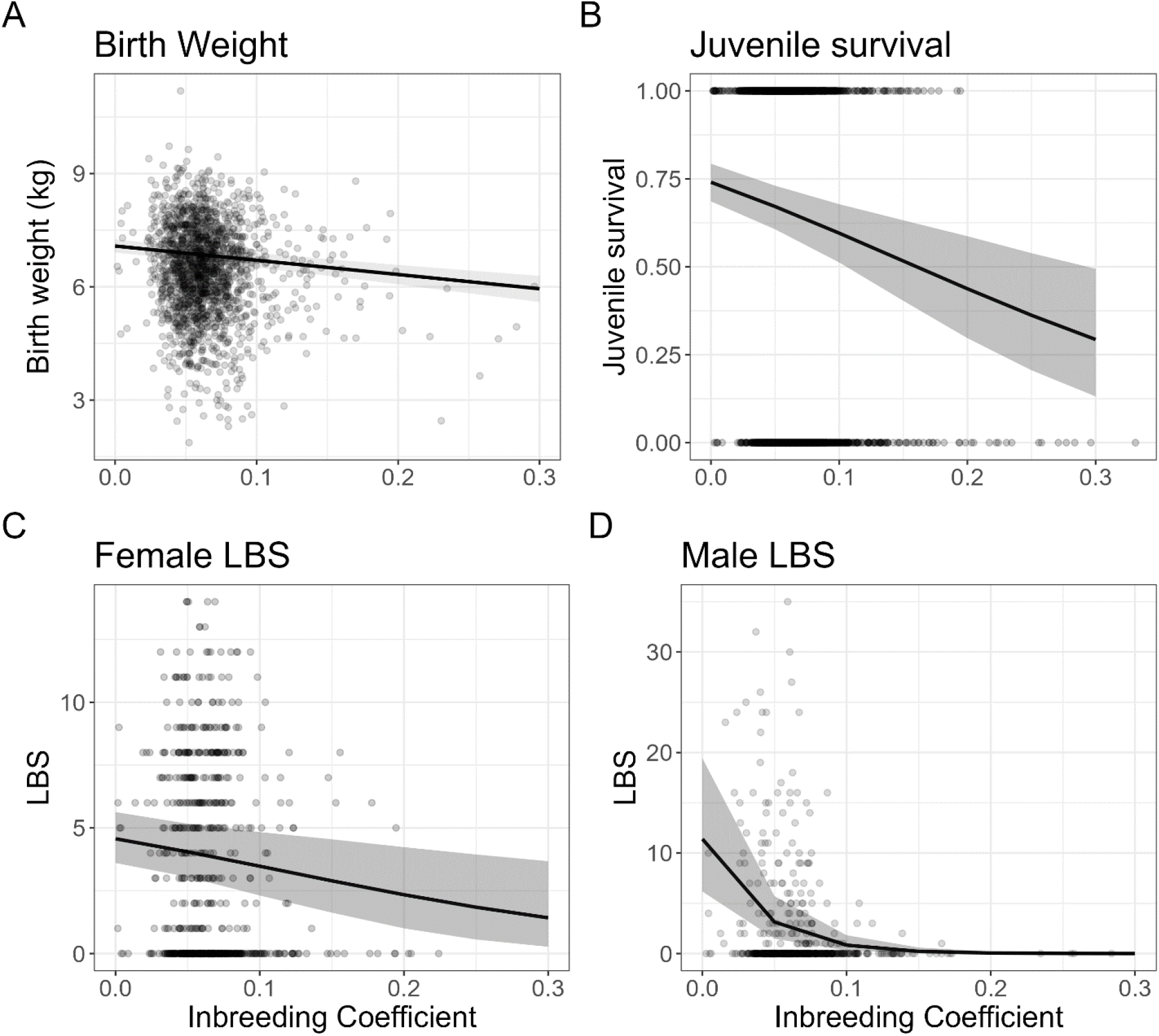
Predicted birth weight (A), juvenile survival probability (B), and lifetime breeding success in females (C) and males (D) using model estimates for increased inbreeding coefficient, where F_ROHChr_ is the same value on all chromosomes. Solid lines shows the mean posterior estimate for an average individual and grey area shows the 95% confidence intervals for the posterior estimate based on stored iterations. For figures C and D predictions use the combined processes of hurdle/zero-inflation and truncated Poisson/Poisson. Points show the raw data. For figure A, capture weights are only shown for individuals <48 hours old at capture for plotting purposes.

### Chromosome-specific effects of inbreeding

Across all four tested traits the majority of chromosomes showed a marginal decrease in fitness with increasing F_ROHchr_ (Fig. 4), but no chromosome had a significantly detrimental effect compared to the average effect of inbreeding (Fig. 3).

**Figure 3.**
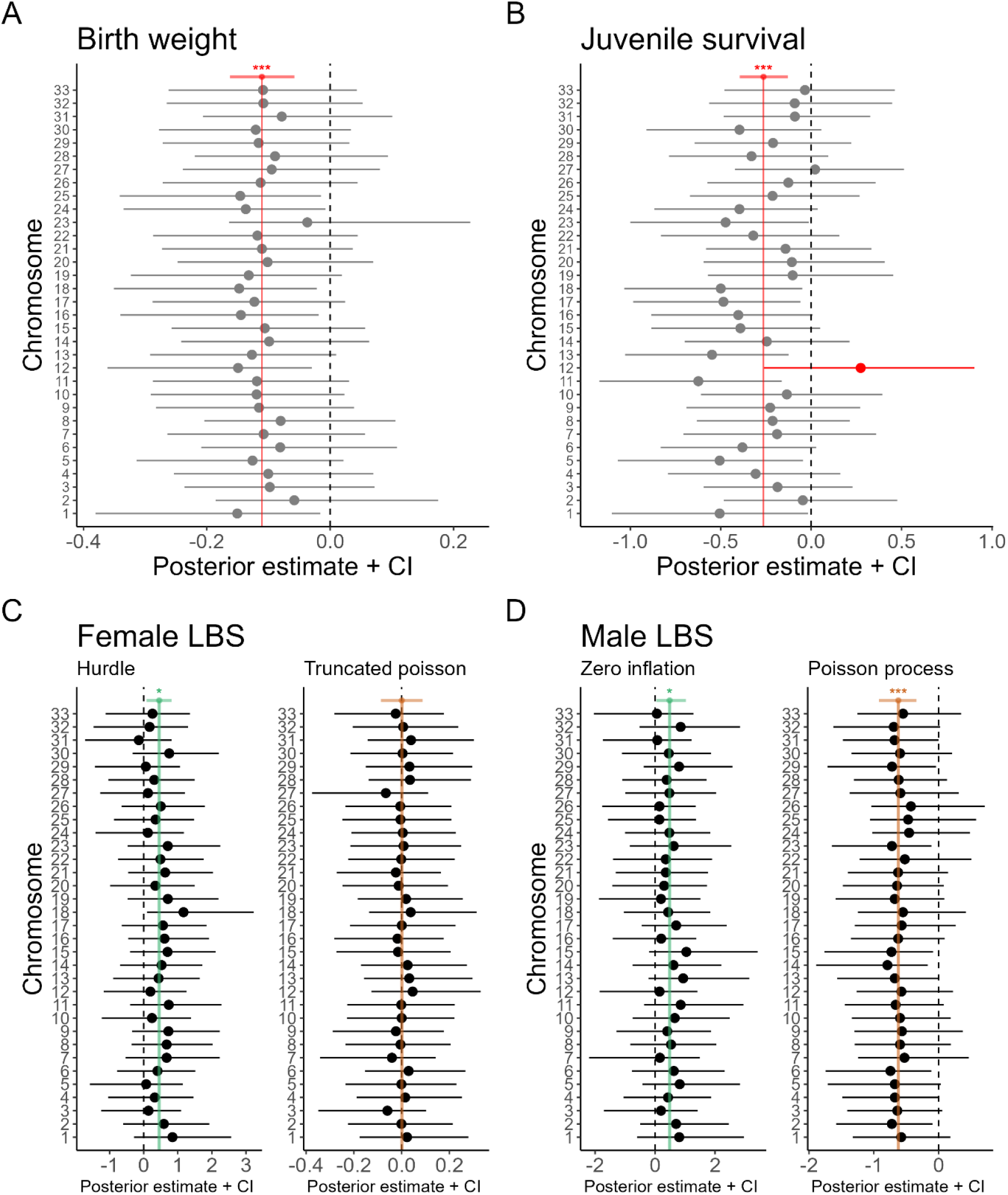
Deviation of chromosome specific inbreeding coefficients from the summed inbreeding effect of all chromosomes for birth weight (A), juvenile survival (B), and lifetime breeding success for females (C) and males (D). Vertical dashed line shows H_0_ (no effect of inbreeding). Figures C and D shows the hurdle/zero-inflation on the left and the truncated Poisson/Poisson process on the right. Vertical solid red line in A and B shows the estimated effect of summed chromosomal inbreeding coefficients on trait. Vertical solid green lines shows the mean estimated effect of summed chromosomal inbreeding coefficients on the hurdle (C) and zero inflation (D) expressed on the logit scale as the probability that LBS=0, where a positive estimate indicates an increase in this probability. Vertical orange lines show the average inbreeding effect on the truncated Poisson (C)/ Poisson (D). Horizontal coloured line shows the 95% confidence intervals for the summed estimate and associated significance value (*: p-value<0.05, ***: p-value<0.001). Grey/black dots show the chromosome-specific point estimates and horizontal lines show the 95% confidence interval. Figure B chromosome 12 is highlighted red as confidence intervals do not overlap the summed effect.

**Figure 4.**
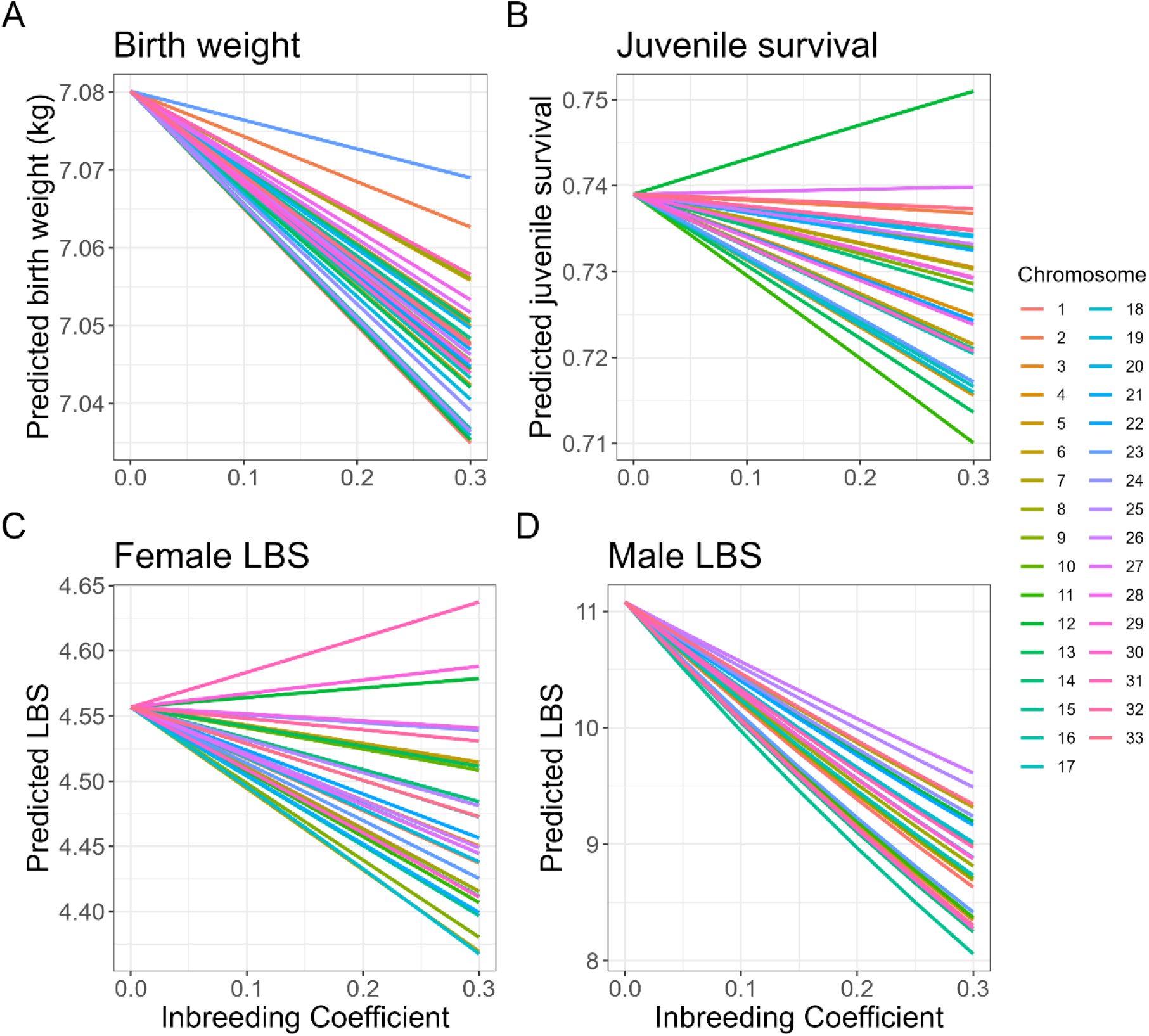
Mean predicted birth weight (A), juvenile survival (B), and lifetime breeding success for females (C) and males (D) for increasing F_ROHChr_ when chromosomes are treated independently in a hypothetical scenario i.e. F_ROHchr_ only increases on the focal chromosome and all other F_ROHchr_ = 0.

When individual chromosome effects are summed they produce the overall inbreeding depression shown in Figure 2. One exception is shown for inbreeding on chromosome 12 (highlighted red in Fig. 3B) which was significantly associated with an increase in survival (although the lower 95% confidence interval nearly touches the average). Increased F_ROHchr_ on chromosome 12 predicts a modest increase in survival rate, whereas for all other chromosomes it predicts a lower rate of survival (Fig 4).

### Effect size and chromosome length

The effect size of each chromosome on birth weight was negatively correlated with the length of the chromosome (Supplementary Figure 2) i.e. larger chromosomes contributed more to inbreeding depression, although this relationship did not cross the significance threshold (p-value: 0.054). A similar trend could be observed for other traits but as above the relationships were not significant (p-value juvenile survival: 0.29, p-values 0.088 and 0.45 for female LBS for hurdle and truncated Poisson respectively, p-values 0.61 and 0.64 for male LBS for zero inflation and Poisson respectively).

### Power of multi-membership approach

The power to detect chromosome-specific inbreeding depression using the multi-membership model approach was dependent on the sample size and the lethality of increasing F_ROHchr_. We simulated juvenile survival data and F_ROHchr_ values using the overall juvenile survival rate and F_ROHchr_ mean and standard deviation representative of our empirical dataset (See supplementary material for details). Greater sample sizes increased the power to detect chromosome-specific inbreeding depression. We also show that increasing the maximum value of F_ROHchr_ which leads to mortality actually decreases the power to detect such effects. For example, supplementary Figure 4 shows that an effect that causes mortality when F_ROHchr_ is greater than 0.4 will only be detected if the sample size is ∼6,000.

## Discussion

The distribution and effect size of loci underpinning inbreeding depression in wild populations using fitness data has seldom been investigated. Our main aim was to gain a better understanding of the genetic architecture of inbreeding depression in the population of red deer inhabiting the Isle of Rum. In order to do so, we estimated the inbreeding effect size on individual chromosomes for traits previously identified as experiencing inbreeding depression. Using a larger sample size, we confirmed previous estimations of inbreeding depression for birth weight, juvenile survival, and lifetime breeding success (Huisman *et al*, 2016). On average, calves produced from a full-sibling or parent-offspring pairing (F_ROH_≈0.25) were ∼12% lighter at birth than a completely outbred calf. We also show calves with an inbreeding coefficient of 0.25 had a ∼38% lower probability of survival compared to outbred calves. Considering the average weight of a calf at capture is ∼7kg, the reduced birth weight in inbred individuals potentially impacts their survival probability, as smaller calves have higher mortality in the first few months of life (Clutton-Brock *et al*, 1982). During refinement of the model of juvenile survival we also fitted birth weight as an explanatory variable but found no difference in the mean effect estimates. As a result, we chose to omit it from the final model as inbreeding depression in birth weight appears to be a component of inbreeding depression in juvenile survival.

We also replicate findings of inbreeding depression in lifetime breeding success (LBS) found in Huisman *et al* (2016), where inbred individuals of both sexes were less likely to have at least one offspring. Given the knowledge of inbreeding depression on juvenile survival this result is perhaps expected. Nearly all females that survive to age three years reproduce (Pemberton *et al*, 2022). Therefore, most females that have no offspring are those that didn’t survive to reproductive age. For males, a large number of individuals do not sire any offspring either because they die before reproductive age or because they are unsuccessful in their reproductive efforts (Pemberton *et al*, 2022). A high proportion of males modelled in the zero-inflation process will then be individuals that do not survive to adulthood. As we have shown, an individual’s inbreeding coefficient affects its likelihood of surviving to reproductive age and hence its likelihood of reproducing. Therefore, the inbreeding depression in LBS is partly influenced by the effects of inbreeding on juvenile survival. We further show that inbred males also had a lower number of total offspring compared to outbred males, analogous to Huisman *et al* (2016). This suggests that inbred male are less competitive. Competition for matings with females is intense in red deer and the fighting success of a male is dependent on his size, weight and condition, which in turn plays a major role in whether he secures a harem and mates successfully (Clutton-Brock *et al*, 1982). Inbred males may be in worse condition, less likely to win fights and therefore unlikely to hold large harems and sire multiple offspring. Alternatively, the lower number of offspring may be caused by the residual effects of juvenile survival. Individuals with an LBS of zero captured in the Poisson process may be a mix of those that do not survive to adulthood and those that achieve no successful matings. Therefore, the effect on inbreeding depression in juvenile survival may be represented in both the zero inflation and the Poisson process.

Although the pedigree-based expectation of identity-by-descent is the same for all chromosomes within an individual, the correlations between chromosome-specific inbreeding coefficients within individuals were relatively low here, due to recombination and independent segregation (Figure 1). This is further consistent with the overall low level of inbreeding in the population and the relatively low estimated identity disequilibrium (Huisman et al, 2016). As a consequence, there is potential for different chromosomes to vary in their inbreeding effects, but at the same time are not entirely independent. Under this reasoning and with the aim of identifying chromosomes with disproportionally large inbreeding depression effects, we estimated the deviation of chromosome-specific inbreeding coefficients from the overall effect of inbreeding using multi-membership models.

Across all investigated traits we found no chromosomes that have a significantly detrimental effect in comparison to the overall effect of inbreeding. Most chromosomes do show a marginal decrease in fitness with increased chromosomal inbreeding, which, when summed results in the observed inbreeding depression. This indicates that in this population there are no chromosomes that contribute disproportionately to the observed inbreeding depression, instead inbreeding depression is the cumulative result of many small effect loci spread across all chromosomes. An alternative explanation for our results is that we did not have the power to detect differences between chromosomes, although, we demonstrate that for a simulated dataset with an equal sample size, chromosome-specific inbreeding that has a lethal effect on survival would be detected using these methods (Supplementary Figure 4). Additionally, we show that the length of the chromosome scaled with the effect of inbreeding depression in some of the tested traits (birth weight and probability of a female having at least one offspring), although these relationships do not cross the significance threshold. Nevertheless, this suggests that longer chromosomes contribute more to inbreeding depression because they contain more mutations, similar to the assumptions of loci with additive causal effects in polygenic traits (Robinson *et al*, 2013; Visscher *et al*, 2007; Yang *et al*, 2011).

If inbreeding depression in the red deer on Rum is caused by many loci with small effects this could partly explain its persistence. The efficiency of purging deleterious loci depends on a number of factors including the selection coefficient and the rate of inbreeding. Mildly deleterious recessive mutations are not exposed to such extreme purifying selection and therefore are not easily purged (Charlesworth and Willis, 2009; Wang *et al*, 1999). In contrast, mutations with high selection coefficients (lethal or extremely detrimental fitness effects) will be quickly purged from populations. Therefore, as inbreeding depression is caused by loci with modest or weak effects in this population, selection may not be effective in removing the contributing alleles although this does not mean that mutations with major deleterious effects have not existed in the past. Rather, the lack of evidence for such mutations implies they may have already been purged from the population.

The population history of the red deer on Rum may further explain the persistence of inbreeding depression. A recent study on wild Alpine Ibex showed that the number of deleterious mutations in a population increased with the strength of a population bottleneck (Grossen *et al*, 2020). Populations that have undergone an extreme population bottleneck will have a fast level of inbreeding (high ΔF), meaning that purifying selection is inefficient and deleterious mutations can accumulate (Charlesworth and Charlesworth, 2010; Reed *et al*, 2003; Wang *et al*, 1999). The red deer population on Rum experienced a minor bottleneck ∼150 years ago when various introductions were made to the island from multiple sources and now has an effective size of ∼175 (Gauzere *et al*, 2022; Marshall, 1998). This population history resulting in a moderate ΔF per generation could have allowed for the purging of strongly deleterious mutations but been inefficient for weakly or moderately deleterious mutations, hence, their effects persist. While the risk of extinction as a result of an accumulation of these mutations, so called mutational meltdown, is a concern for populations with an effective size less than 100 (Lynch *et al*, 1995), the Isle of Rum deer population exceeds this number. Despite this, studies on mutational meltdown are scarce in wild populations (Robinson *et al*, 2023), so it will be useful to monitor the population, particularly its effective size, in future.

It is important to note that increased statistical power (through increasing the number of individuals and/or using denser genomic markers) would probably uncover loci with contributions to inbreeding depression which is further supported by our simulations (See supplementary Figure 4 and supplementary material). As is the case with complex traits, increasing the number of loci and individuals usually increases the number of loci associated with the trait, each explaining a small amount of variance (Visscher *et al*, 2010; Yengo *et al*, 2022). Indeed, one recent study of a wild population with more individuals, denser SNPs, and higher individual inbreeding coefficients detected large effect loci contributing to inbreeding depression (Stoffel *et al*, 2021). Increasing power in this way is obviously difficult in studies of wild populations and still does not guarantee the statistical power to detect loci with major deleterious effects. If only a small number of individuals have a ROH at a locus, the power to detect the effect would be insufficient, even if it was lethal.

Such loci would be expected to occur at low frequencies, hence, the power to detect them will be low as there are few individuals carrying the causal allele (Kardos *et al*, 2016). However, in smaller populations with higher average relatedness, deleterious recessive alleles could reach high frequencies due to drift, consequently increasing the power to detect them (Kardos *et al*, 2016). Therefore, the conclusions we make surrounding inbreeding depression being due to many loci with modest/weak effects spread genome-wide would likely remain, even with an increase in power.

One major exception to the general pattern was found on chromosome 12. Increased inbreeding on this chromosome was associated with significantly increased juvenile survival. For example, if an individual had a chromosomal inbreeding coefficient of 0.3 on chromosome 12 and was completely outbred on all other chromosomes their survival probability would increase by 1.2% compared to if the individual was outbred on all chromosomes (Fig. 4B). Interestingly, chromosome 12 is also associated with an increase in female LBS, likely due to the effects of survival on LBS, but this association is not statistically significant (Fig. 4C). This may represent an example of a beneficial recessive allele which when homozygous confers a survival advantage. In livestock species such regions are often associated with economically important genes where being homozygous can result in an increased yield, hence they are usually under intense artificial selection (He *et al*, 2020). However, in this population, increased inbreeding on chromosome 12 will incur higher inbreeding genome-wide, resulting in an overall depression in fitness.

In summary, we show evidence for ongoing inbreeding depression in birth weight, juvenile survival and lifetime breeding success in a wild population of red deer. Most chromosomes show a minor decrease in fitness with increased chromosome-specific inbreeding coefficients but, except for one instance discussed above, none are significantly different from the average effect of inbreeding. Therefore, in this population inbreeding depression is probably the result of many loci each with a modest or weak effect. Theory predicts that such mutations will be inefficiently purged and can lead to the persistence of inbreeding depression.

## Acknowledgements

We thank NatureScot for permission to work on the Isle of Rum. We thank the many field workers that have helped to collect samples and life histories on Rum, particularly Alison and Sean Morris. We are also grateful to Jarrod Hadfield, Joel Pick, Kirsty Macphie and Albert Phillimore for statistical advice and discussions. We also thank Martin Stoffel for discussions as well as Ino Curick and Simon Martin for discussions and comments on an earlier version of the manuscript. The long-term project and this research were funded by the UK Natural Environment Research Council, and most of the SNP genotyping was supported by a European Research Council Advanced Grant to J. M. Pemberton. A. Hewett was supported by an E4 NERC DTP studentship (NE/S007407/1). S.E. Johnston is supported by a Royal Society University Research Fellowship (UF150448). Genotyping was conducted at the Wellcome Trust Clinical Research Facility Genetics Core. We are grateful to the Darwin Tree of Life project for the physical map positions used.

## Author contribution statement

J.M.P coordinated sampling and genotyping and supervised the fieldwork. A.M.H conducted analyses and drafted the manuscript. All authors contributed to revisions.

## Data availability

Data will be archived upon acceptance.

