## Supplementary figures and tables for "Genetic architecture of inbreeding depression may explain its persistence in a population of wild red deer"

### *Supplementary material*

To simulate the effect size detected by the multi-membership approach we generated $F_{ROHchr}$ values for 33 chromosomes using a random generation of numbers from a truncated normal distribution using a mean $F_{ROHchr}$ of 0.065 and a standard deviation of 0.2, where values were truncated between 0 and 1. We also simulated survival values using a random generation of binomial distribution using the probability of survival as 0.58. We generated these numbers for 6 different sample sizes: 150, 500, 1500, 3000, 4500 and 6000. To simulate a 'lethal mutation' on a chromosome we altered the survival of individuals with $F_{ROHchr\_5}$ (inbreeding coefficient on chromosome 5) greater than a certain value and scored them a survival of 0, we did this for 5 values of $F_{ROHchr}$: 0.1, 0.2, 0.3, 0.4 and 0.5. For example, for value 0.1 every individual with $F_{ROHchr} > 0.1$ scores a survival of 0. Multi-membership models on the simulated datasets were then run in MCMCglmm using the default iteration settings. The estimated effect size and associated 95% confidence intervals for the chromosome of interest were extracted and compared between simulations. See supplementary Figure 4 for results.

### *Supplementary Figures*

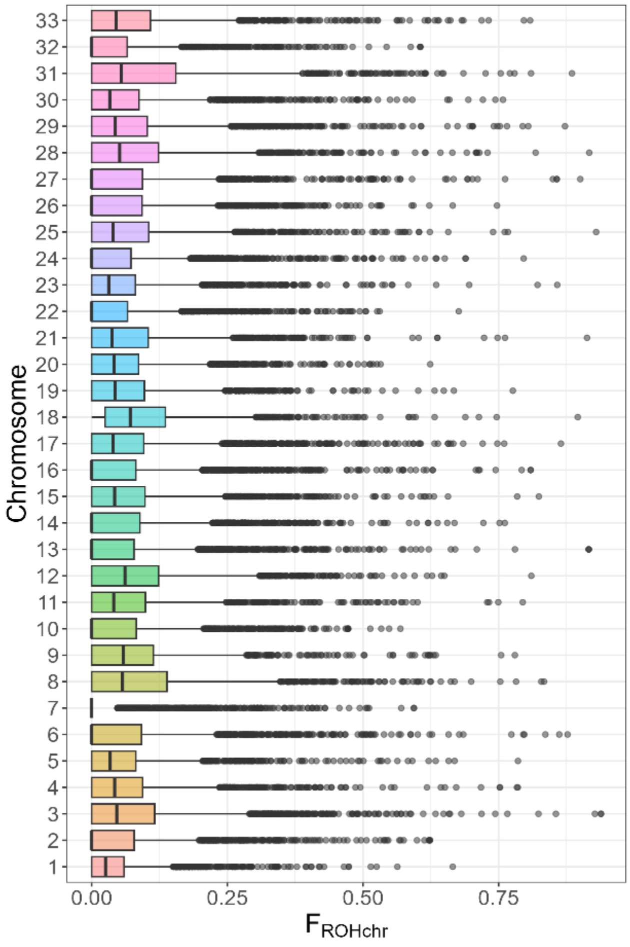


**Supplementary Figure 1** – Box plots showing chromosome-specific inbreeding coefficients ($F_{ROHchr}$) for all 3198 individuals on all 33 autosomes. Mean $F_{ROHCh}$r for each chromosome shown by the solid black line within the box, outliers shown as dots.

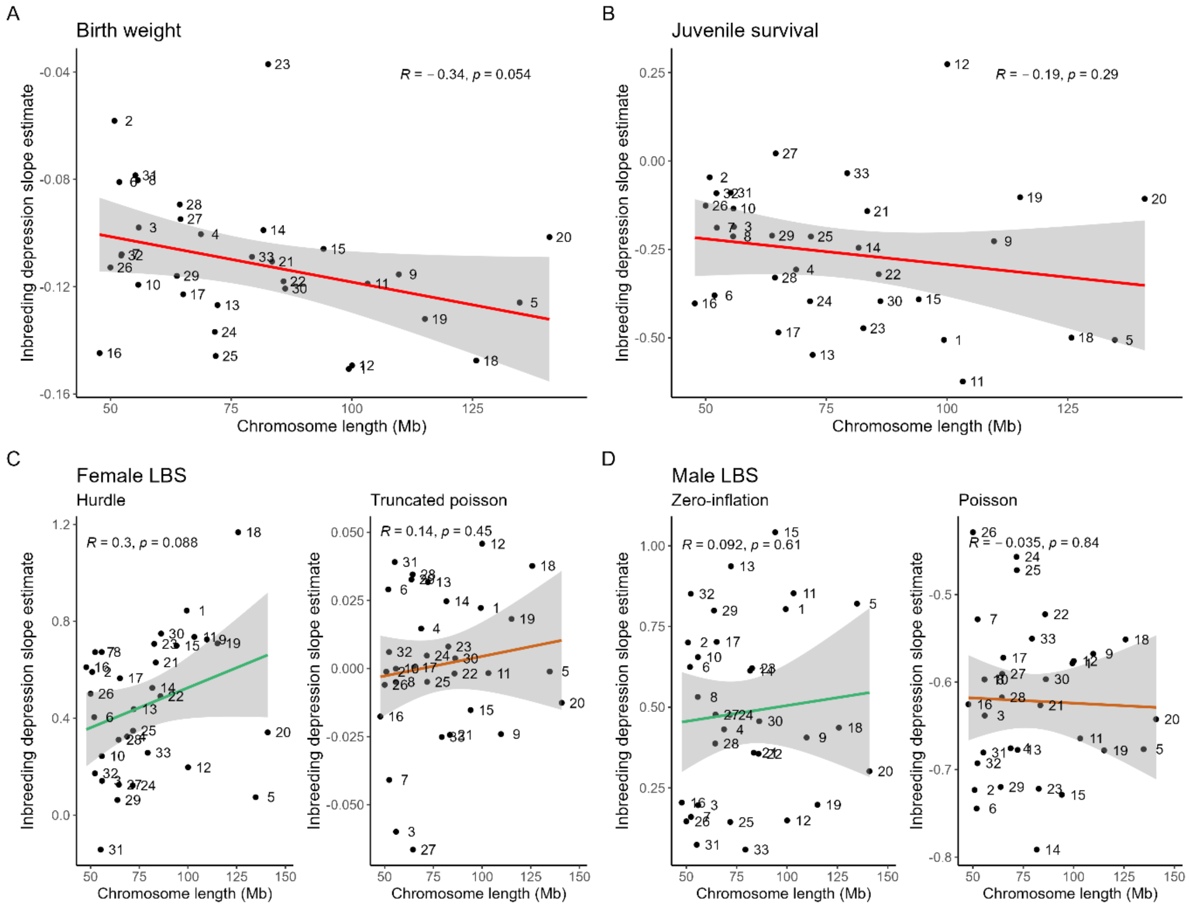


**Supplementary Figure 2** - Inbreeding depression slope estimate plotted against chromosome length in Mb for birth weight (A), juvenile survival probability (B), and lifetime breeding success in females (C) and males (D), line shows the slope of the relationship as a linear model. Figures C and D show hurdle/zero-inflation (expressed as the probability that LBS=0, where a positive relationship shows inbreeding depression increases with chromosome size) and truncated Poisson/Poisson separately. Labels show chromosome number. Top shows estimate of the correlation and associated p-value

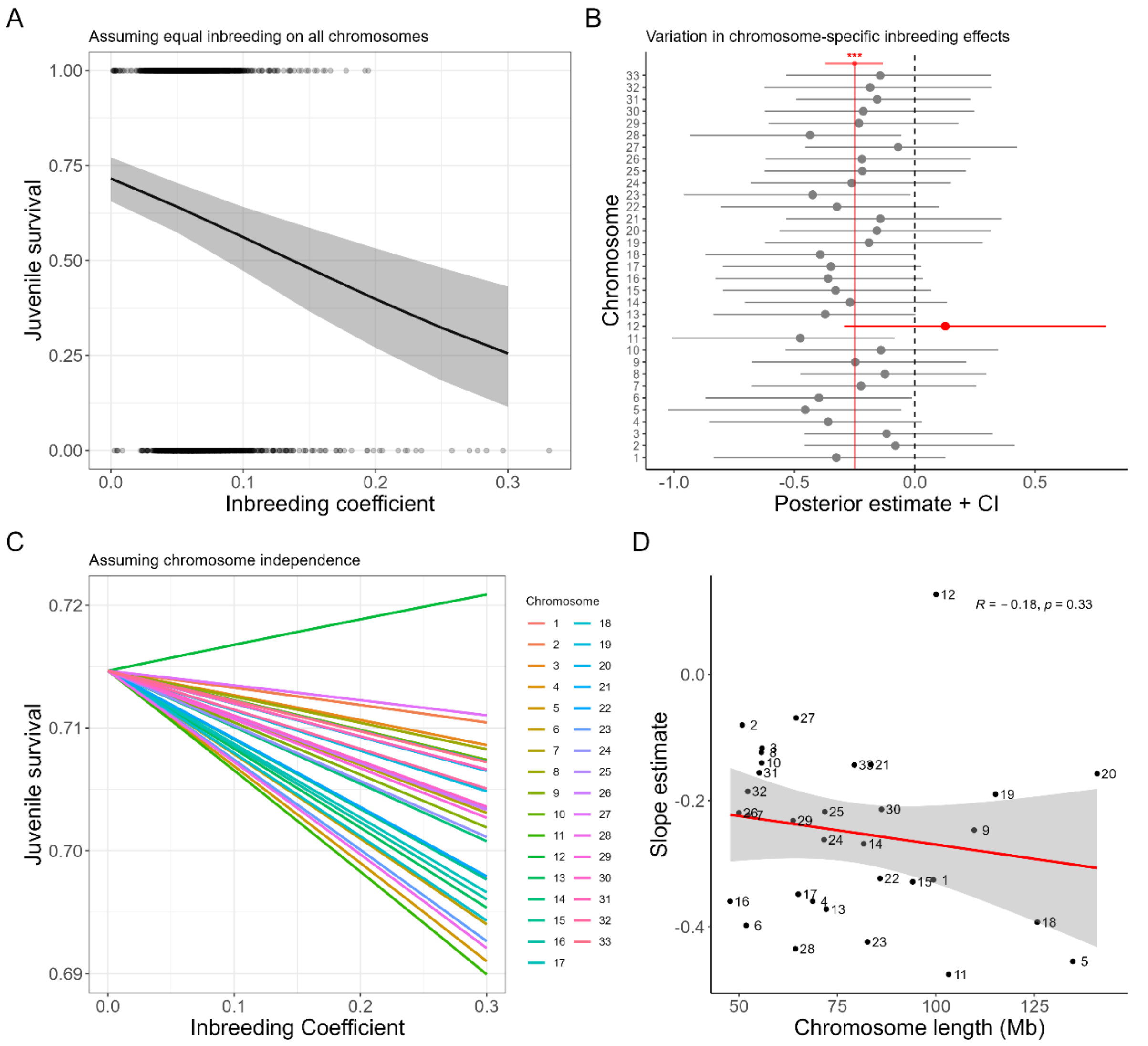


**Supplementary Figure 3** - Repeat of Figures 1B (A), 2B (B), 3B (C)and supplementary Fig2B (D) with the addition of birth weight fitted as a fixed effect in the model of juvenile survival.

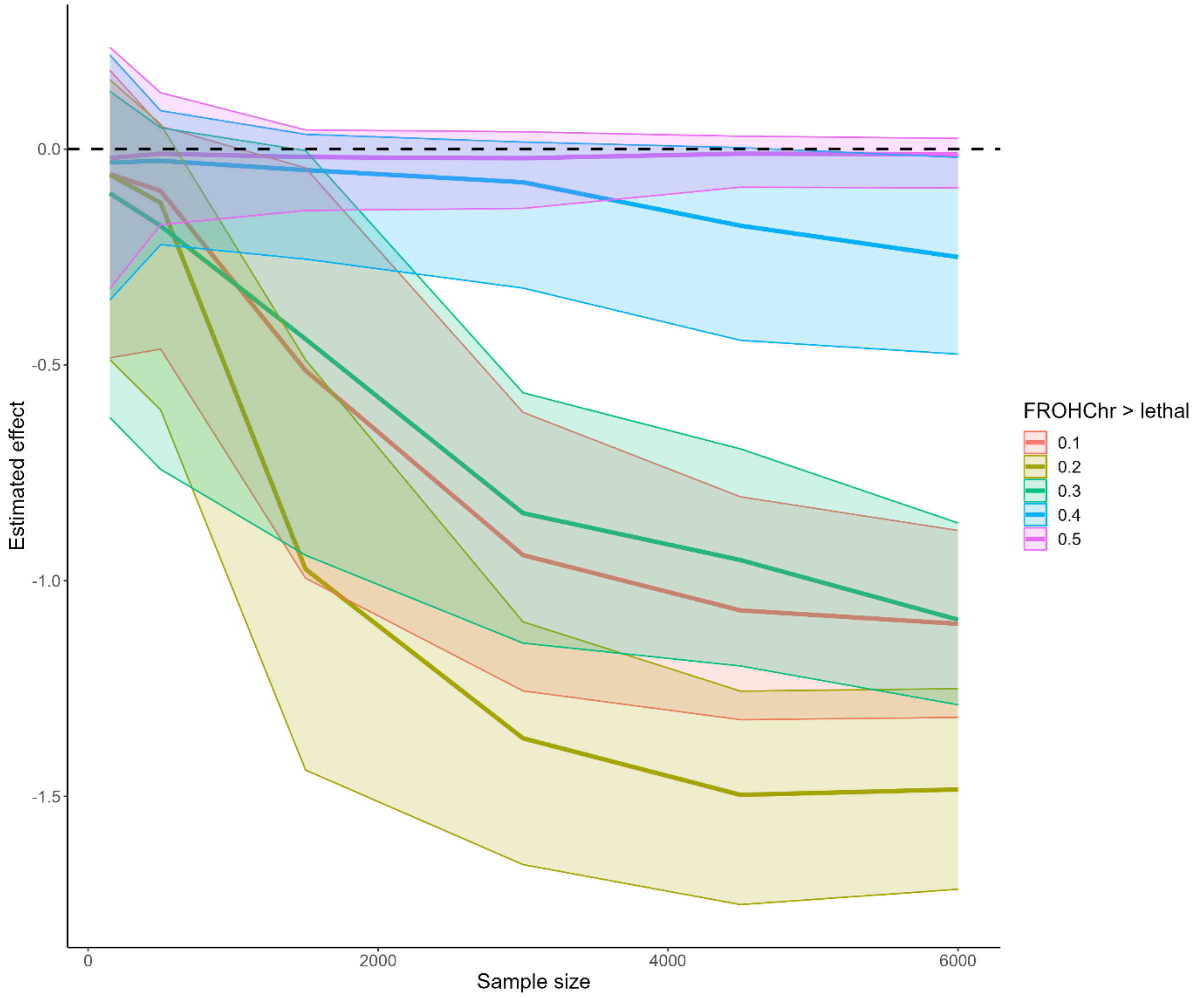


**Supplementary Figure 4** - Estimated effect of a chromosome conferring lethality in simulated data using the multi-membership model approach with increasing sample size. Confidence intervals that overlap the black horizontal line shows that no effect would be detected. Colours represent $F_{ROHchr}$ values that confer lethality e.g. red shows a simulation where every individual with $F_{ROHchr} > 0.1$ scored a survival of 0.
